# Habitat fragmentation causes coevolutionary burning spots

**DOI:** 10.1101/116293

**Authors:** H. De Kort, M. Baguette, J.G Prunier, M. Tessier, J. Monsimet, C. Turlure, V.M. Stevens

## Abstract

Habitat fragmentation increasingly threatens the services provided by natural communities and ecosystem worldwide. An understanding of the underlying eco-evolutionary processes in natural settings is lacking, yet critical to realistic and sustainable conservation. Through integrating the multivariate genetic, biotic and abiotic facets of a natural community network experiencing various degrees of habitat fragmentation, we provide unique insights into the processes underlying community functioning in real, natural conditions. The focal community network comprises a parasitic butterfly of conservation concern, and its two obligatory host species, a plant and an ant. We show that fragmentation of butterfly habitat has the potential to impair the balance between dispersal and coevolution. This process can cause coevolutionary burning spots of decreased genetic diversity and therefore of increased extinction risk. We stress that ignoring such eco-evolutionary feed-backs inherent to the very functioning of natural communities can strongly impact their persistence.

**One Sentence Summary:** Communities under threat of habitat fragmentation suffer increased extinction risk through coevolutionary overheating.

---

Biotic interaction networks including pollination, competition and parasitism play a key role in community functioning and persistence, but global environmental changes compromise their ecological and evolutionary stability (*1–3*). Fueled by ongoing land conversions, habitat fragmentation in particular increasingly degrades ecological communities and ecosystems across the globe (*4, 5*). Yet attempts to mitigate these impacts are hampered by a markedly poor understanding of the eco-evolutionary dynamics driving biotic interaction networks under threat of habitat fragmentation (*6–8*). Indeed, owing to technical challenges associated with studying eco-evolutionary community processes in natural settings, non-experimental empirical studies demonstrating how biotic interactions implicate the ability of species to withstand habitat fragmentation are lacking.

Theoretical models and experimental work on microbial communities predict that habitat fragmentation and community functioning are linked through the effects of landscape connectivity on dispersal, the latter defining individual movements across the landscape (9–*12*). According to these *in silico* and *in vitro* studies, intermediate levels of dispersal facilitate species evolution and species sorting concordant with local biotic and abiotic conditions, whereas low (or high) levels of dispersal can deplete (or swamp) locally adapted communities. However, the exact processes determining eco-evolutionary outcomes are highly variable, shaped by species’ genetic architecture, their phenotypic variation, the strength and amount of interactions with local biotic and abiotic variables, and the eco-evolutionary feedbacks resulting from interactions between each of these components (*7, 13*). This complexity is compromising empirical validation of theoretical predictions, urging for a more realistic perspective on the impacts of altered landscape connectivity on communities and ecosystems (*14*–*16*).

Analogously to the interwoven nature of community dynamics and adaptive evolution, we propose the integration of landscape genetic disciplines in community ecology (*17*) to address this complexity *in situ.* Ecological and evolutionary processes including drift, gene flow, and abiotic and coevolutionary selection directly influence patterns of genetic variation within and among communities. Coevolution in particular, i.e. the process by which two or more interacting species reciprocally affect each other’s evolution, may give rise to specific genetic signatures when communities are spatially structured. More specifically, spatial community structure can generate a geographic mosaic of coevolutionary hot spots featured by strong reciprocal selection among interacting species (*18–20*). Where local coevolution prevents successful immigration of maladapted individuals, or when it favours reduced dispersal ability in isolated habitats (*21*), such selection mosaics can result in paralleled genetic structure among interacting species (*22*). Assessing population genetic variation within and among the different species of a community network may therefore shed light on the impact of habitat fragmentation on community dynamics.

Here we present the first empirical data showing how habitat fragmentation can impact the eco-evolutionary dynamics propelling communities in natural settings. We focus on a community system highly appreciated for its conservation value: the threatened and specialized European butterfly *Phengaris* (*=Maculinea*) *alcon alcon* (the Alcon blue, hereafter “butterfly”) and its two obligatory hosts, a rare grassland plant *Gentiana pneumonanthe* (the march Gentian, hereafter “plant”) and an ant of the genus *Myrmica* (here the common *M. scabrinodis,* hereafter “ant”) (Fig. 1A) (*23*, *24*). Due to habitat fragmentation, the butterfly faces local extinctions across its range, even where its hosts remain locally abundant (*25*). We focus on a mountainous landscape in the French department Ariège (4890 km^2^), where we investigated all known butterfly communities. These community sites are spatially structured into four disjoint clusters (metacommunities), with among-cluster distances (>10km) exceeding the known maximum dispersal distance of the butterfly (0.5km) (Fig. 1B-D, Fig. S1). The communities span an altitudinal gradient from 400 to 1017 m, with patch sizes varying between 275 and 17,359 m^2^ (Table S1).

**Fig. 1.**
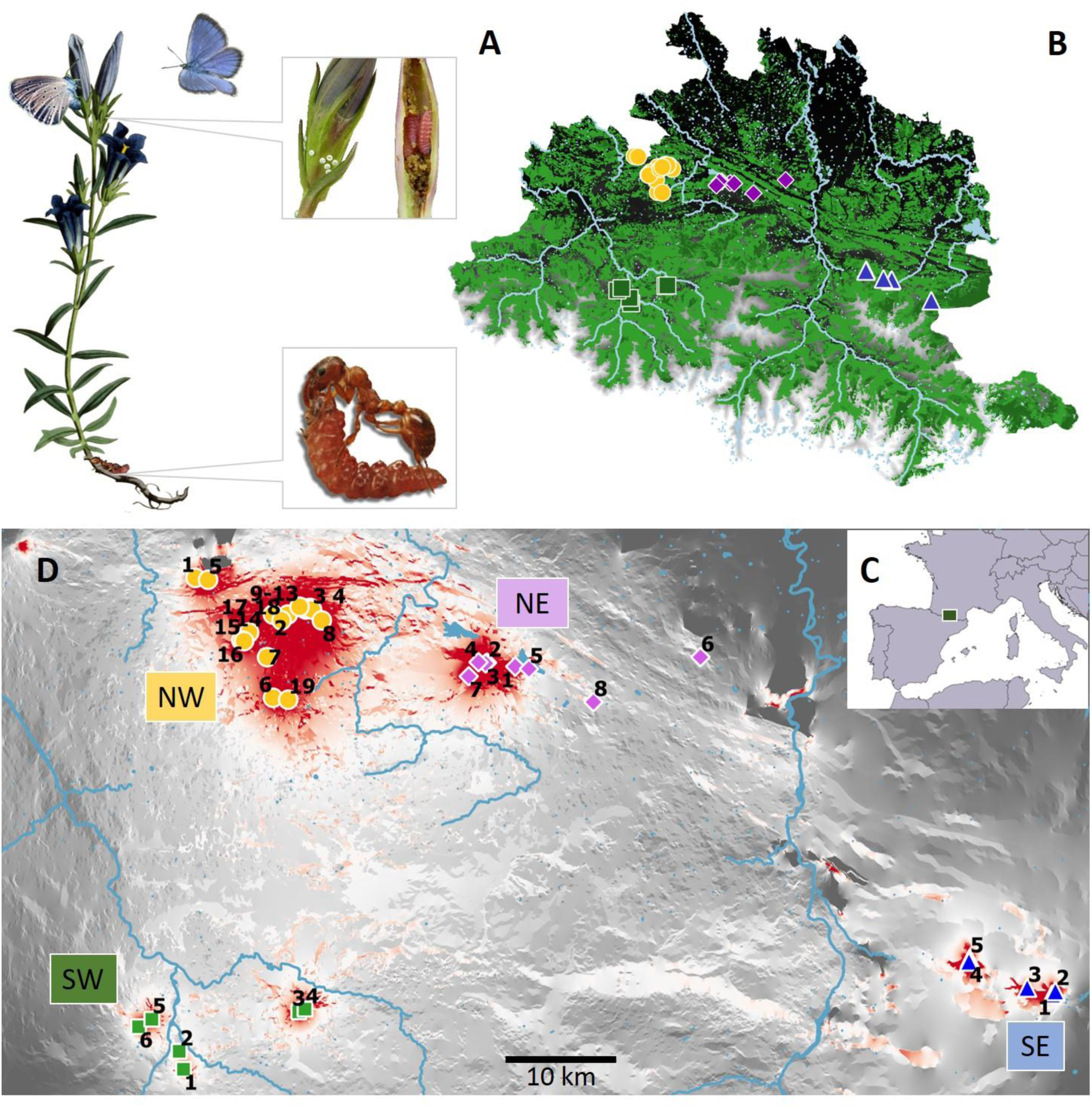
Study system. **A**. Butterfly community. After mating, females Alcon blue butterflies (*Phengaris [=Maculinea] alcon*) lay their white eggs onto gentian (*Gentiana pneumonanthe)* flower buds. Small caterpillars develop into the bud, at the expense of gentian’s ovules. After their 3^rd^ molt, the caterpillars leave the plant and are adopted by ants (*Myrmica scabrinodis*), that rear them into their underground nest in preference to their own brood. This social parasitism ends after metamorphosis. For visibility, ant size was exaggerated relative to plant size. **B**. Overview of the study landscape in which the samples were collected, with tree cover shown in green, and with a black to white gradient reflecting altitude. Shape and colour of sampling locations correspond to their geographical cluster (yellow circles: NW, purple diamonds: NE, blue triangles: SE, green squares: SW). **C**. Positioning of sampling area in the French central Pyrenees (SW Europe). **D**. Landscape connectivity map of the study area representing butterfly dispersal probability. Red/grey colours represent habitat favouring/impeding dispersal. Rivers (blue) align with the main valleys. Geographical and demographic details about the sampling sites can be found in Table S1.

Based on land use, plant requirements, and the biology of the butterfly, we mapped butterfly dispersal probability across the landscape to obtain a relevant measure of landscape connectivity for this species (Fig. 1D, Table S2-3). Because habitat fragmentation deteriorates landscape connectivity through reductions in the amount of habitat suitable for dispersal and breeding, and through increased geographical distance among breeding sites, it typically disrupts the genetic integrity of butterfly populations (*26*). In line with this unfavourable trend, we found that decreased landscape connectivity increased the genetic distance between the butterfly populations under study (Fig.S2). This finding is consistent with field observations obtained during daily individual butterfly movement monitoring, showing some dispersal within geographical clusters (3.55% butterflies recaptured in another site vs. 35.49% recaptured on site), limited dispersal among the nearest clusters (0.51%), and no dispersal among the more distant clusters (Fig. S1). By hindering butterfly movements across the landscape to various extents, habitat fragmentation thus resulted in a spatial mosaic of highly and poorly connected sites (Fig. 1D).

We then explored how the genetic diversity within communities was affected by landscape connectivity, local habitat size and altitude. Genetic diversity was calculated for 22 butterfly, 37 plant and 29 ant populations, based on high-quality single nucleotide polymorphisms obtained through pooled RADseq (Fig.S3). We found a marked negative effect of decreasing landscape connectivity on butterfly genetic diversity (Fig.2A, Fig.S4, Table S3), implying a harmful effect of habitat fragmentation on its population dynamics. We also observed that both butterfly and plant genetic diversity decreased with altitude (Fig.2B). Because census population sizes did not follow an altitudinal pattern (Fig. S5), we devote this result to historical post-glacial migration to higher altitudes followed by population expansion. Together, landscape connectivity and altitude explained 74.23% (R^2^_adj_) of the variation in butterfly genetic diversity.

**Fig. 2.**
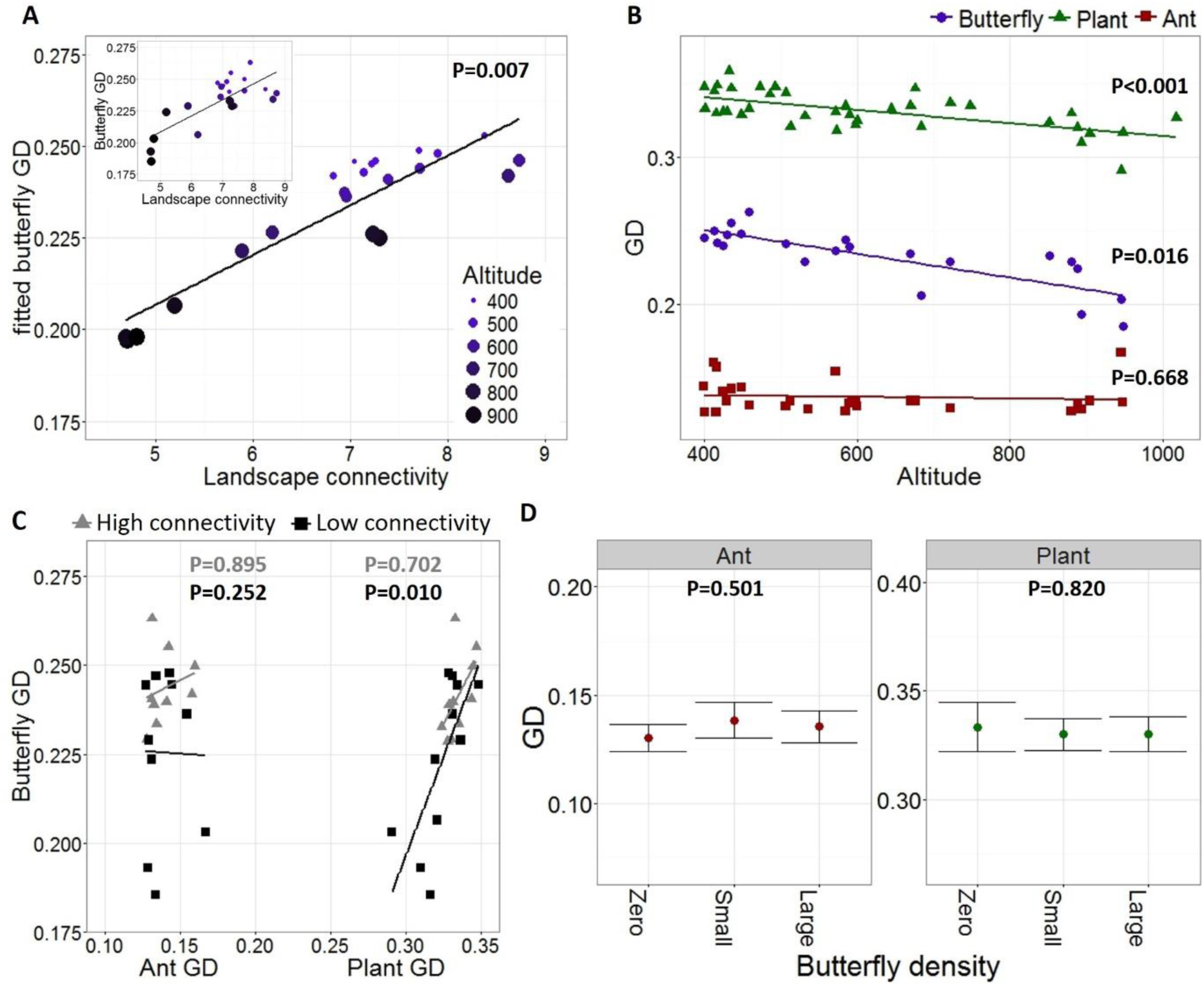
Genetic diversity (GD) patterns of the butterfly community network. A. Correlation between landscape connectivity and fitted and original butterfly H_E_ values. Fitted values result from a weighted linear model with landscape resistance, altitude and patch size (Log10-transformed) as explanatory variables (Table S4). Dots get darker and larger with altitude. **B**. Correlation between altitude and genetic diversity of each species. **C**. Relation between butterfly and host genetic diversity for both high landscape connectivity (>7.2, the median) and low landscape connectivity (<7.2). **D**. Parasitic effect of the butterfly (based on local population densities) on genetic diversity of hosts. Zero butterfly density refers to recently extinct populations.

Given the absence of a negative correlation between butterfly and host genetic diversity, our results also suggest limited parasitic impact of the butterfly on its two host species. To corroborate this theory, we compared ant and plant genetic diversity between sites with high vs. low butterfly population density (i.e. high vs. low parasitic pressure), and sites where the butterfly got extinct in the last 5 years (Table S1). The genetic diversity of the hosts did not decrease with increased butterfly density (Fig.2D), indicating that the hosts are not markedly impacted by butterfly population dynamics. Importantly, this also suggests that conservation efforts aiming to increase connectivity for butterflies would maintain the functioning of the community network without compromising the host species.

The strong positive correlation between butterfly and plant genetic diversity, but not between butterfly and ant genetic diversity (Fig.2C, Table S4), corresponds to the shared effect of altitude on their genetic diversity (Fig.2B). However, because the correlation between butterfly and plant diversity is significant only where landscape connectivity is relatively low (Fig.2C), genetic diversity losses in the plant (the least mobile partner of the community) may have negative consequences for butterfly genetic diversity where habitat fragmentation impedes butterfly gene flow between populations. To confirm the genetic dependence between the butterfly and the hosts in the context of habitat fragmentation, an understanding of both the abiotic and coevolutionary drivers of community dynamics is required.

We therefore investigated the contributions of abiotic landscape variation to butterfly and host genetic structure, using multivariate landscape and community genetic approaches (Table S4). To approximate the abiotic (incl. spatial) environment, we modelled latitude, longitude, altitude, habitat size and landscape connectivity (for the butterfly) or geographical isolation (for the hosts). We thus also accounted for all existing environmental clines that are generally known to covary with these factors (e.g. temperature and precipitation). The abiotic variables explained a significant proportion of the community genetic structure, varying from 13.72% for ants over 28.03% for plants to 32.95% for butterflies. Latitude, longitude and altitude all contributed to the notable genetic structure of the butterfly and the plant (Fig.3A-C, Fig.S6). The genetic structure of the ant was less pronounced, although the southern clusters were strongly differentiated owing to geographical isolation and altitudinal differences (Fig.3C). Together, these findings indicate an important role for spatial structure and the associated abiotic clines in driving genetic differences among communities. On top of the observed spatial effects, landscape connectivity influenced butterfly genetic structure (Fig.3A), suggesting that habitat fragmentation interferes with its metapopulation dynamics. As expected from the complexity inherent to natural communities, a large part of the genetic structure remains unexplained, and could be partially due to biotic interactions which are most often ignored in landscape genetic studies.

**Fig. 3.**
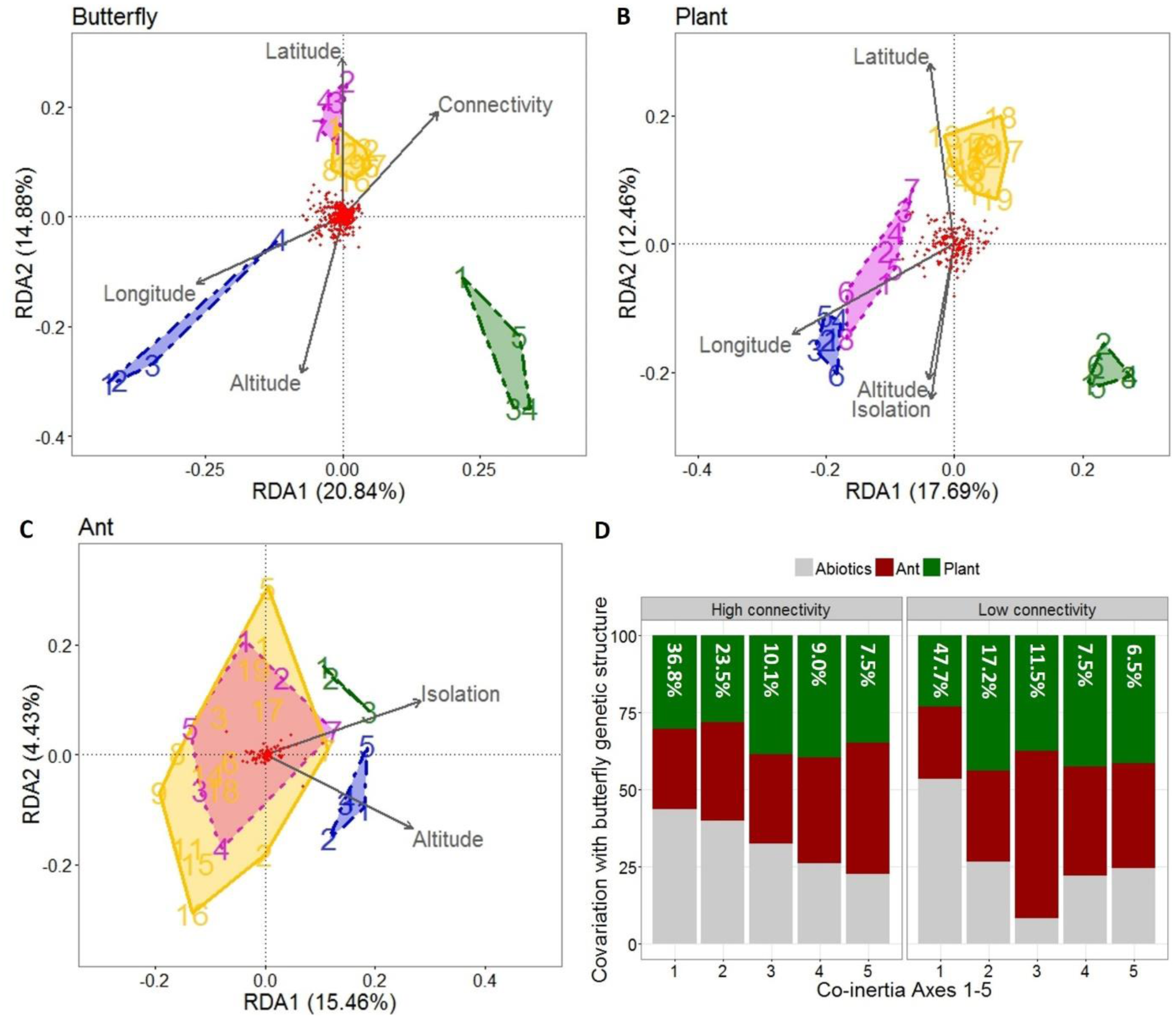
Role of environmental factors contributing to the genetic structure of the butterfly community network. **A-C**.Triplots representing the genetic structure of butterfly, plant and ant, respectively, and the abiotic factors significantly contributing to this structure (Table S4). Colours correspond to spatial structure in Fig. 1D. **D**. Contribution of host genetic structure and abiotic factors to the genetic structure of Alcon populations. The first five axes of the co-inertia analyses explained a total of 86.89% and 90.34% of the covariation between the Alcon genetic matrix and the environmental matrix (altitude, longitude, latitude, landscape connectivity, ant and plant genetics), for the highly and poorly connected populations, respectively (Table S4). Contributions of each axis to the total coinertia are shown in white at the top of the bar plot.

We thus incorporated the genetic structure of the host species as a proxy for the biotic environment of the butterfly, to identify the proportion of its genetic structure covarying with host genetic structure while accounting for the abiotic environment. The resulting model (Table S4) provides insights in unique, independent processes driving butterfly genetic structure, captured by so-called co-inertia axes, in addition to shared, interdependent processes. To assess the effect of habitat fragmentation on the degree of covariation (i.e. co-inertia) between butterfly and host genetic structure, we contrasted between poorly and highly connected butterfly populations. Substantial variation in butterfly genetic structure can be explained by host genetic structure and abiotic variation, although this covariation was only significant for the poorly connected sites (covariation coefficient = 0.85^ns^ and 0.84** for high and low connectivity, respectively, Table S4). Overall, most of the butterfly genetic variation co-varied with the abiotic and biotic environment simultaneously (Fig.3D), indicating shared effects between the abiotic variables and the host genetic structure on butterfly genetic structure. This is in line with our observation that shared abiotic conditions drive the genetic structure of the three species (Fig.3A-C).

Where landscape connectivity is high for the butterfly (Fig.3D, left panel), we observed strong common contributions of abiotic variables and host genetic structure to butterfly genetic structure (equivalent contributions to co-inertia axes 1-3). In addition, a limited part of the butterfly genetic structure can be uniquely explained by host genetic structure, as shown by the disproportional contribution of plant genetic structure to co-inertia axis 4, and of ant genetic structure to axis 5. Because this parallel genetic structure between the butterfly and its two hosts cannot be fully explained by abiotic variables, we suspect that reciprocal coevolutionary selection between the butterfly and its two hosts has reduced effective gene flow between the communities, thereby synchronizing the genetic structure of the host-parasite network. This notion is concordant with previously documented coevolutionary shifts in flower phenology (*G. pneumonanthe* plants) and surface chemistry (*Myrmica* ants) to escape local infestation by *P. alcon* butterflies (*23, 24).* Where postponed plant flowering and altered ant surface chemistry prevent reproduction of desynchronized butterfly immigrants, local coevolution has the potential to increase the genetic distance between communities, resulting in parallel genetic structure among the interacting species. Correspondingly, the genetic structure of another *Myrmica* ant species has been demonstrated to be closely associated with its local surface chemistry signature, evolved to counter butterfly impacts (*23*). We conclude that spatial community structure provides opportunities for coevolution, while frequent local butterfly immigration (under high landscape connectivity) maintains relatively high levels of genetic diversity within metapopulations.

Where landscape connectivity is low (Fig.3D, right panel, Fig.4B), a more profound relation between butterfly and host genetic structure was apparent. Host genetic structure (plant: axes 2; ant: axis 3) and abiotic factors (axis 1) independently covaried with butterfly genetic structure (Fig. 3D), likely reflecting intense coevolutionary genetic parallelism. This result is consistent with the idea that habitat fragmentation can reinforce restrictions on effective gene flow driven by coevolution. Alternatively, coevolution has been shown to favour decreased dispersal in isolated patches as a strategy to maintain local coadaptation (*21*), and this could also generate genetic parallelism. However, considering that even the most isolated populations of this study system are potentially connected to others (Fig.1D), selection for reduced dispersal seems unlikely. Because habitat fragmentation intensifies coevolutionary pressures (Fig.3D, right panel), while decreasing genetic diversity (Fig.2A), our findings suggest the existence of coevolutionary “burning” spots, featured by increased local butterfly extinction risk. From a global viewpoint, we argue that the widespread occurrence of coevolutionary interactions renders communities even more vulnerable to habitat fragmentation than generally acknowledged.

**Fig. 4.**
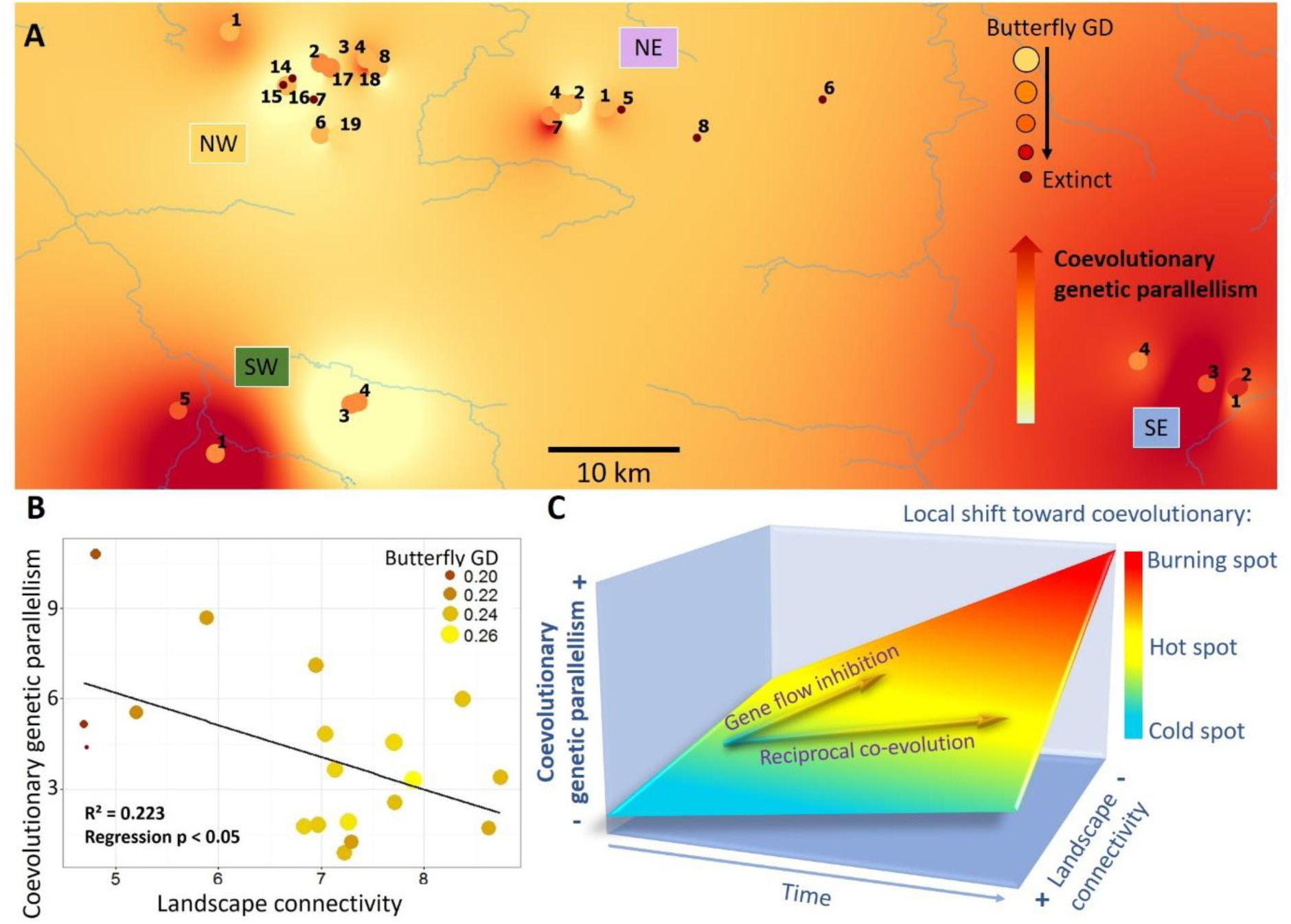
Coevolutionary dynamics over space and time. **A**. Map showing the degree of coevolutionary genetic parallelism between the butterfly and its two hosts. The degree of genetic parallelism likely due to coevolution was derived from a coinertia analysis, through extracting the axis that was mainly represented by host genetic structure (>90%) rather than by abiotic variation (Table S5). Size and colour of site symbols reflects butterfly genetic diversity. Burning spots are featured by a combination of low genetic diversity and high coevolutionary genetic parallelism. The extinctions in the NW cluster are due to recently abandoned grazing management. **B**. Relation between coevolutionary genetic parallelism and landscape connectivity. **C**. Graphical representation of a landscape varying in the degree of connectivity. Communities in overly connected landscapes experience a gene flow overload, resulting in sub-optimal conditions for coevolution (blue, cold spots). The absence of coevolution is mirrored by a lack of coevolutionary genetic parallelism among species. Intermediate landscape connectivity facilitates reciprocal coevolution, giving rise to coevolutionary hot spots (yellow), featured by species that show genetic similarities due to coevolution. Highly fragmented landscapes (poor landscape connectivity) compromise coevolutinary dynamics through selection for decreased dispersal and/or constraints on effective gene flow, finally resulting in coevolutionary burning spots of increased extinction risk.

Our study does not capture causal relationships between landscape connectivity and community dynamics, but it does provide unique perspectives on the strength of *in situ* species’ associations under threat of habitat fragmentation. Decreased landscape connectivity due to land conversion and degradation has the potential to alter the balance between dispersal and coevolution while decreasing genetic diversity, resulting in locally overheated coevolution featured by increased extinction risk (Fig.4). Reconnecting these communities could restore coevolutionary dynamics and simultaneously improve butterfly genetic diversity and population persistence rate. Interestingly, relatively dense butterfly populations with high genetic diversity did not leave a noticeable mark on host genetic diversity. The gradual and reciprocal accumulation of coevolutionary genetic signatures over time likely has allowed coexistence without strong fluctuations in parasite and host genetic diversity. Building on this finding, we emphasize the strength of community genetic studies in detecting eco-evolutionary signatures that have been shaped over contemporary time periods. To shed light on the global extent of our findings, it would be insightful to deploy the used integrative strategy, combining pooled sequencing and multivariate genetic approaches, in other natural settings. Although a pooled sequencing approach complicates the generation of genetic marker-specific inferences (Fig. S3), it offers major advantages for studying *in situ* community dynamics. Its cost-efficiency could allow genotyping hundreds of populations with modest research budgets, and multivariate techniques subsequently allow disentangling the relative contributions of abiotic and coevolutionary drivers of community dynamics.

Predicting how species will cope with global environmental changes is a key ambition in evolutionary and ecological sciences. However, the lack of eco-evolutionary integration in forecasting models may compromise their ability to predict range dynamics and to provide realistic guidelines for sustainable conservation (*27, 28*). Based on our findings, we stress that ignoring the impacts of widespread coevolutionary networks on the ability of species to cope with global environmental changes can considerably affect outcomes of conservation strategies and forecasting models. Moreover, whereas habitat fragmentation has the potential to alter coevolutionary relations irrespective of other global environmental threats, the combined impacts of multiple global environmental changes on community dynamics are expected to be distressing, but have yet to be assessed.

## Acknowledgements

Raw reads are archived at the National Center for Biotechnology Information (BioProject to complete). SNP allele frequencies and flanking sequences are archived at Dryad (to complete).

We were pleased to obtain insightful comments on the manuscript from experts in the field of community ecology and landscape genomics, including Luc De Meester, Michael Hochberg, José MT Montoya, Olivier Rey, Stephanie Manel and Viktoriia Radchuk.

We thank Gaëlle Blanvillain, Sophie Dardenne, and many students for field assistance, and Murielle Richard for lab assistance. Annie Ouin (INRA), Thomas Houet (CNRS), and the Parc Naturel Régional Pyréénes Ariégeoises (Yannick Barascud and Julien Aït El Mekki) provided the GIS resources used for mapping levels of habitat fragmentation. Primary financial support was provided by the ANR GEMS & INDHET (ANR-13-JSV7-0010-01 and ANR-12 -BSV7-0023-02). HDK, VMS, JP and MB are members of the Excellence Lab TULIP (ANR-10-LABX-41).

Hanne De Kort wrote the manuscript, assisted with sampling, and performed the lab work and data analyses. Virginie Stevens and Michel Baguette provided the context of the project and supervised writing, sampling and data analysis. Jérôme Prunier performed the bio-informatics, and Marc Tessier located the community sites and introduced the study system. Camille Turlure assisted with the analysis of the field surveys, and Jérémy Monsimet assisted with sampling. All co-authors also commented on the manuscript.

## Supplementary Materials

Material and Methods

Figures S1-S6

Tables S1-S5

References (29-48)

**Fig. S1.**
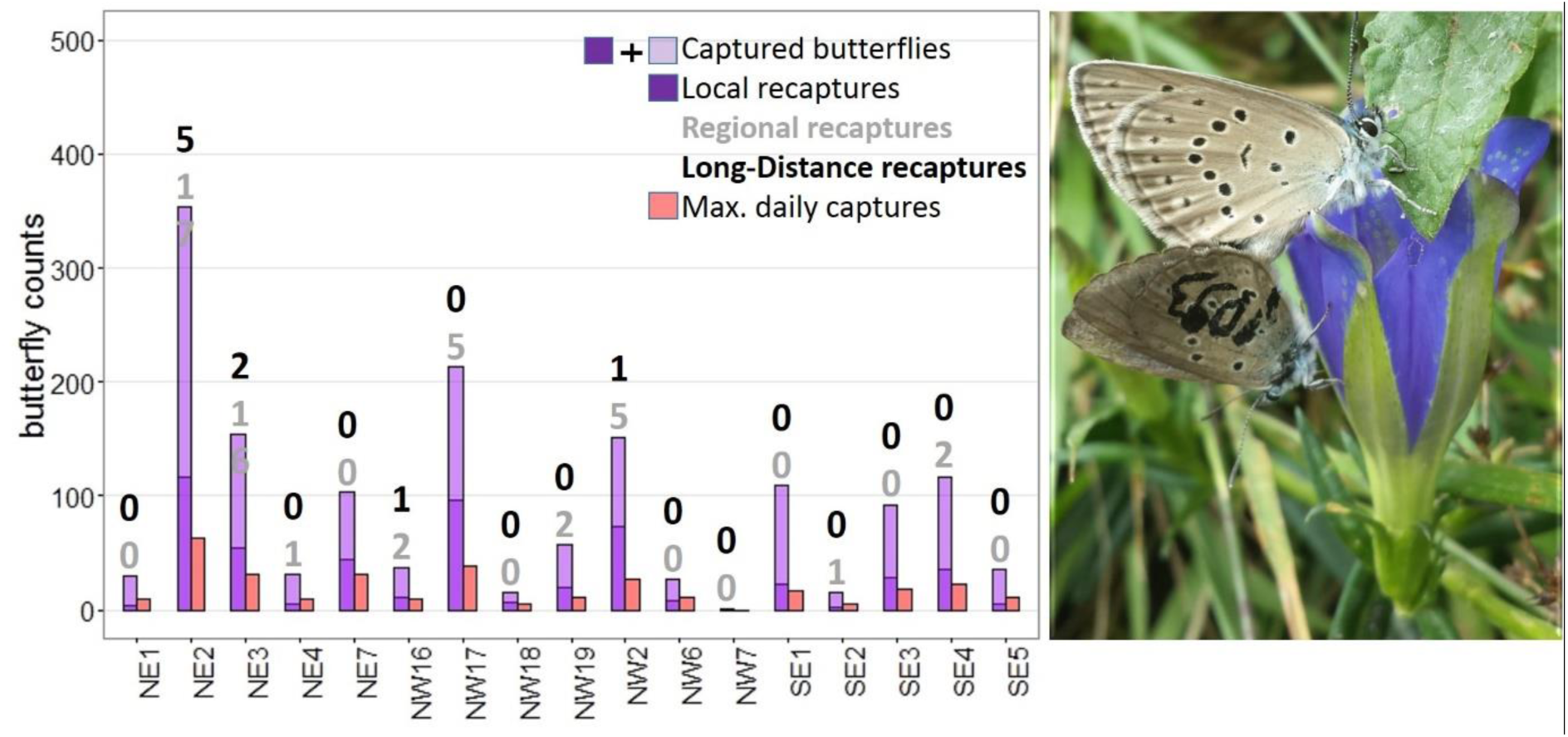
Summary of individual movement monitoring of the NE, NW and SE cluster (summer of 2014), showing number of (i) captured butterflies that were subsequently marked (captured butterflies), (ii) butterflies that were recaptured on site (local recaptures), (iii) butterflies that were recaptured and originated from another site within the same geographical cluster (regional recaptures), (iv) butterflies that were recaptured and originated from another site of another geographical group (long-distance recaptures). Long-distance recaptures only occurred between the NW and the NE metapopulation. These are the first field observations of *Maculinea alcon* dispersal exceeding 0.5 km (*49*). In addition, the number of captures on the best day (day with most captures) is shown. A photograph taken during individual movement monitoring shows a marked butterfly mating with an unmarked butterfly (marked later that day).

**Fig. S2.**
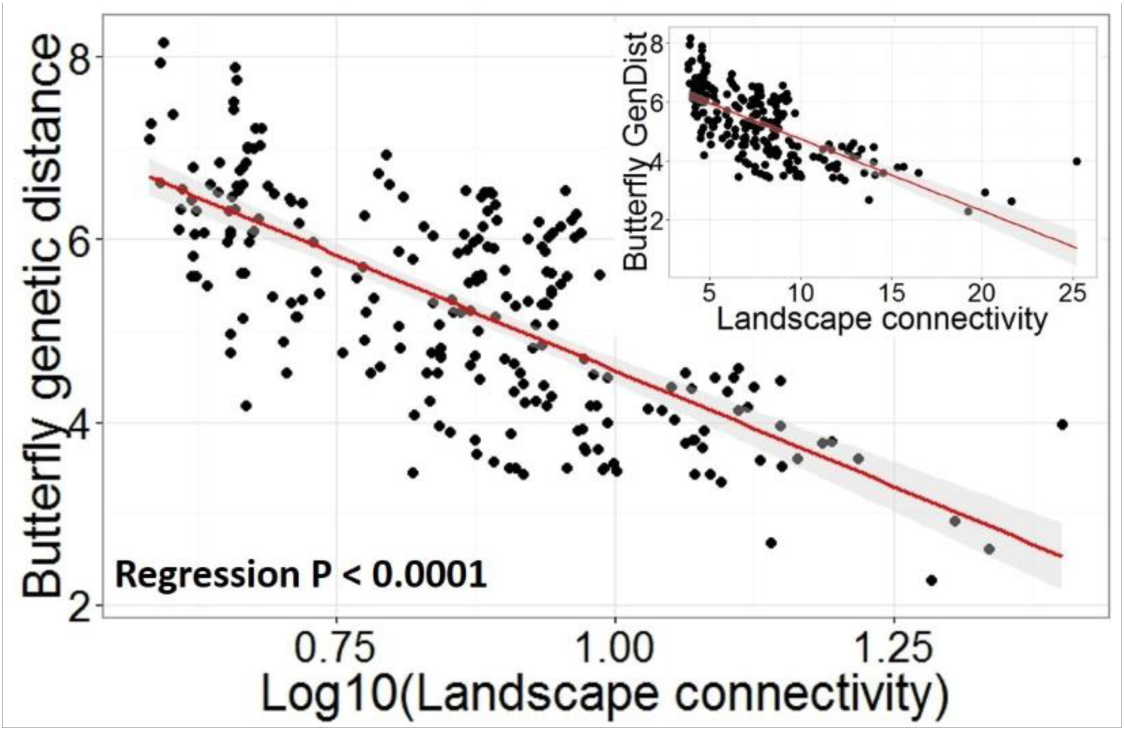
Effect of landscape connectivity on butterfly genetic distance. The scatterplot shows pairwise landscape connectivity values vs. genetic distance. Regression p-value < 10^−16^ for bot log10- transformed and original pairwise connectivity distances. Mantel test results can be found in Table S3.

**Fig. S3.**
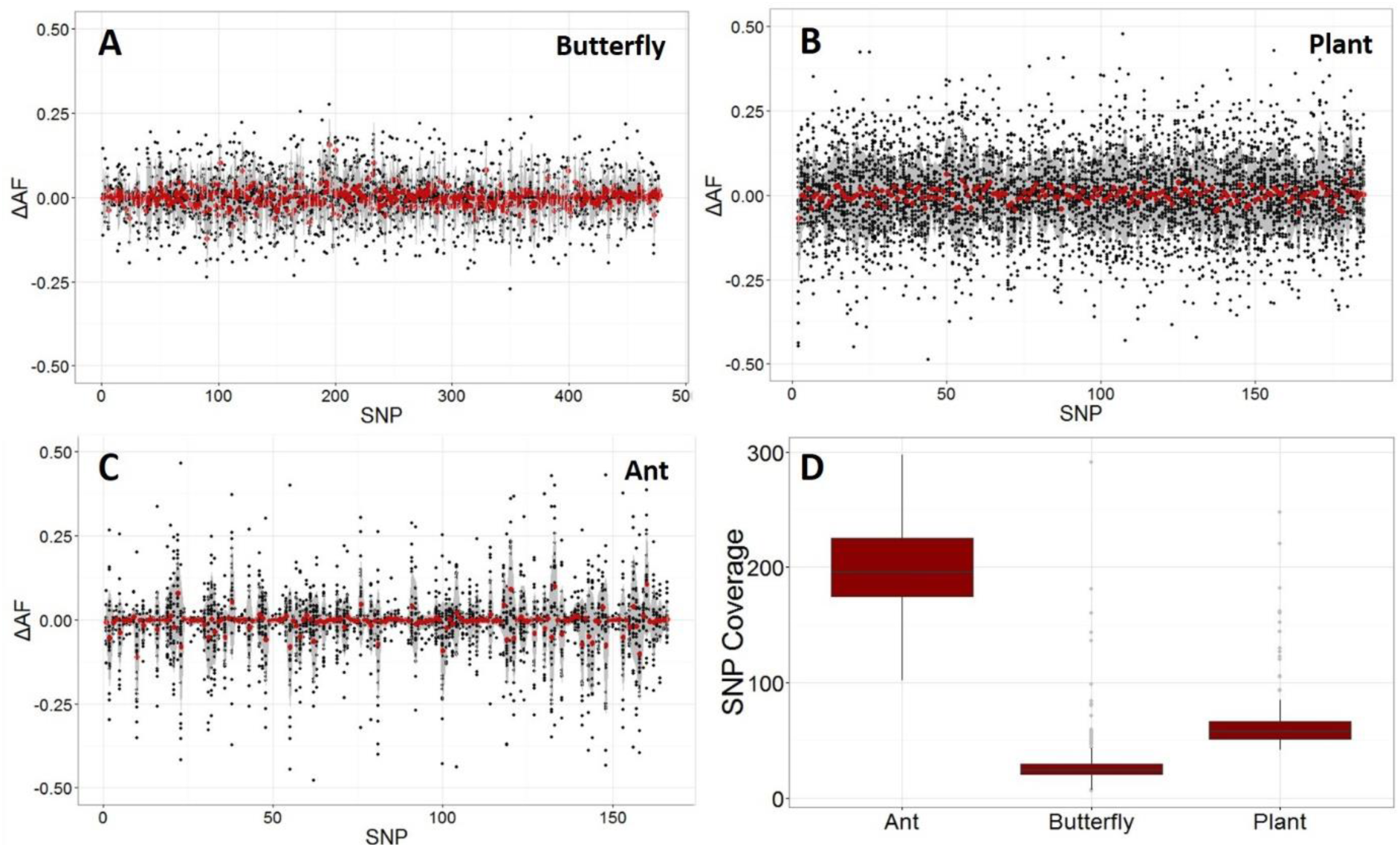
Reliability (**A-C**) and coverage (**D**) of the SNPs used for all analyses. To assess the reliability of the SNP allele frequency (AF) estimates, a total of 6 (butterfly), 30 (plant) and 23 pool replicates (anta) were available, and SNP AFs were compared across pool replicates by calculating the difference in AF (ΔAF) for each SNP in each pair of pool replicates (black dots). Average ΔAF values per SNP are represented by red dots. ΔAF varied substantially, yet randomly, across SNPs and pools, but average ΔAF was <0.01 for each species, rendering the AF estimates suitable for multivariate genetic comparisons among populations and species. A detailed account of the pooled RADseq procedure that resulted in these SNPs is provided in the Material & Methods section.

**Fig. S4.**
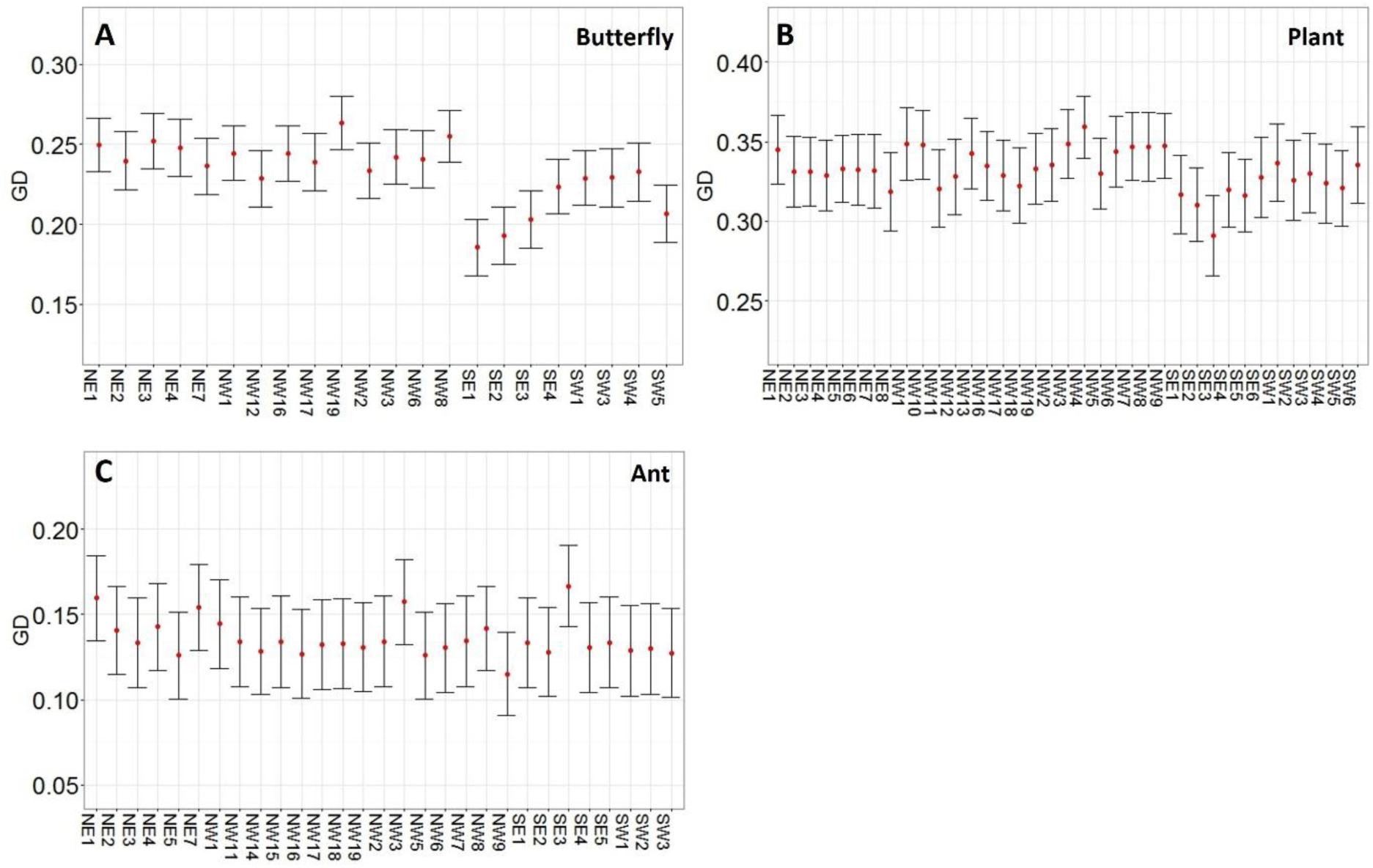
Genetic diversity (mean expected heterozygosity, Table S1) of all successfully genotyped butterfly (A), plant (B) and ant (C) populations.

**Fig. S5.**
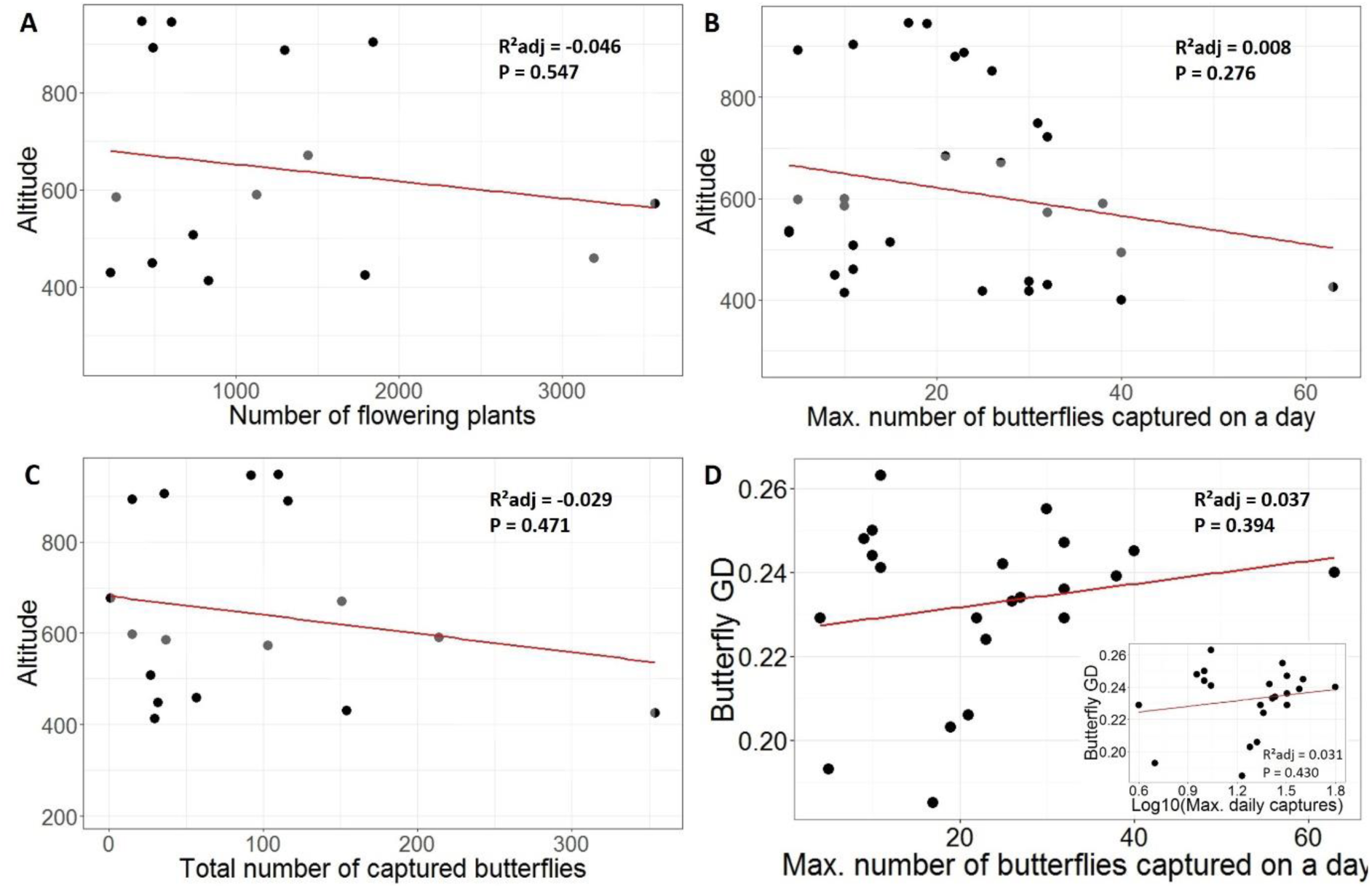
Field summary statistics. **A.** Absence of a relation between plant population size (approximated by flowering plant counts) and altitude. **B-C.** Absence of a relation between butterfly population sizes, based on individual movement monitoring, and altitude. Field methodology is provided in Table S1. Statistics are based on linear regression. **D**. Absence of a relation between butterfly population size and butterfly GD.

**Fig. S6.**
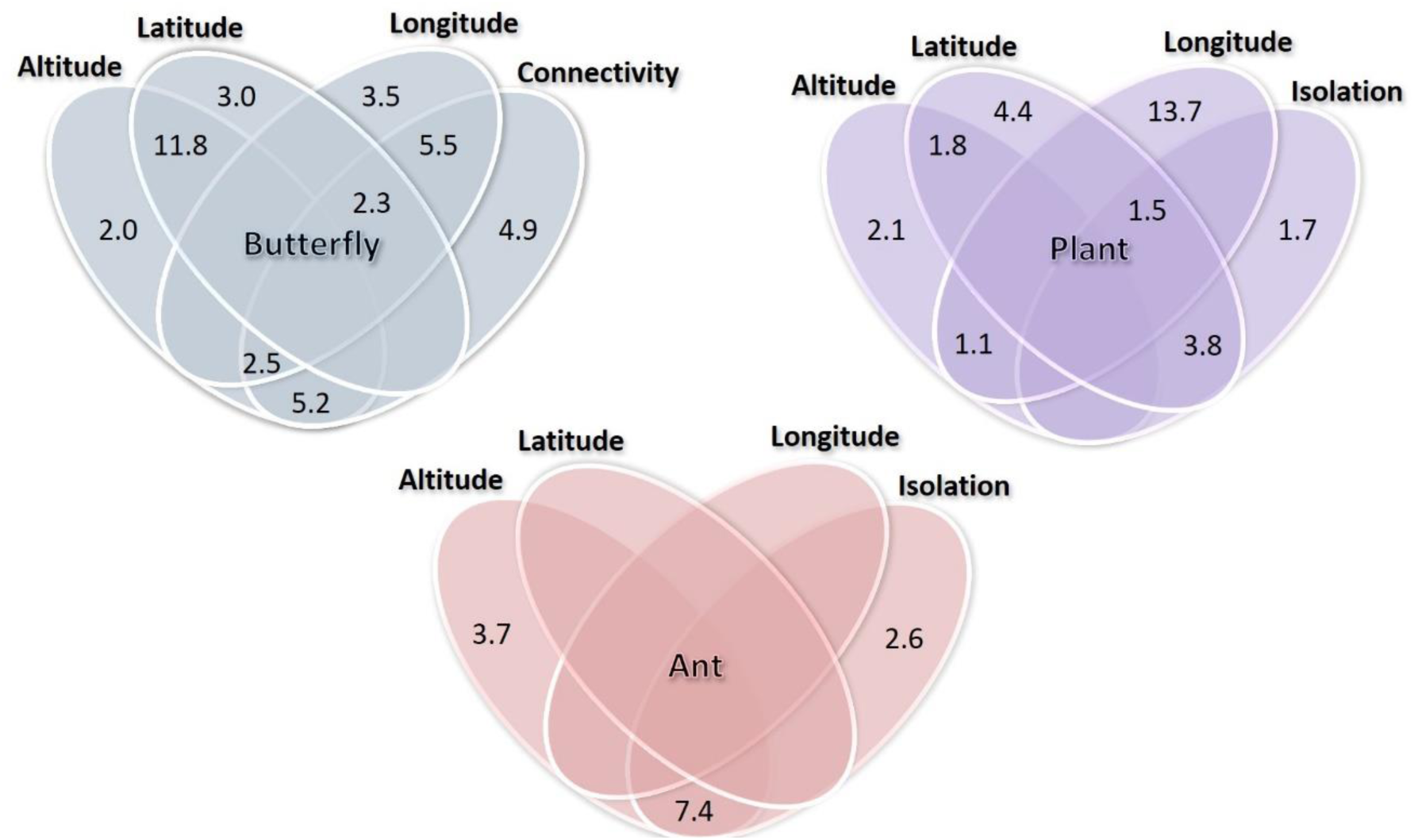
Venn diagram showing unique and common abiotic contributions (%) to butterfly, plant and ant genetic structure. Values were obtained through redundancy analyses with variation partitioning (see Table S4). Only significant contributions are shown.

**Table S1.** Spatial, genetic and demographic population characteristics.

**Table S2.** Assignment of butterfly dispersal costs to (i) land use map categories, (ii) plant presence, and (iii) geological structures that have been found to be associated with plant presence. Higher dispersal costs correspond to lower landscape connectivity. The combination of cost assignments that best explained the genetic distance among butterfly populations (based on partial Mantel tests) is clarified in Table S3. These final dispersal costs were used to calculate pairwise and average landscape connectivity.

**Table S3.** Table of connectivity scenarios with partial Mantel test results. Connectivity scenarios reflect combinations of dispersal costs assigned to land use, plant presence and geology (=likely plant presence).

**Table S4.** Statistical modelling underlying the p-values provided in the text. Models are ranked according to the location of corresponding results in the manuscript. Butterfly habitat size was Log10- transformed to meet model assumptions. To assess the effect of pool size on regression model robustness, regression models were weighted by pool size and compared to unweighted models. Weighing by pool size did not affect model parameters.

**Table S5.** Overall coinertia analysis used to calculate degree of coevolutionary genetic parallellism between the butterfly and its two hosts.

